# GENETIC, MORPHOLOGICAL, AND PHYTOCHEMICAL DIFFERENTIATION OF “CARNAVAL” - A NEW HOP CULTIVAR ADAPTED TO TROPICAL ENVIRONMENT

**DOI:** 10.1101/2025.06.17.660195

**Authors:** Ricardo Beolchi de Souza Lima, Clarissa Hamaio Okino Delgado, Max Vincent Raffaele, Vivian Polotto Croce, José Baldin Pinheiro, Márcia Ortiz Mayo Marques, Jonathan Andre Morales Marroquin, Guilherme Augusto da Silva Martin, Fabíola Ribeiro de Oliveira, Luís Thadeo Poianas Silva, Mateus Mondin

## Abstract

The global hops market generates approximately US$8 billion annually, primarily driven by beer production, the world’s most consumed beverage. Despite this global demand, hop cultivation remains concentrated in temperate regions. Brazil, the third-largest beer consumer, imports over 99% of its hops. This study aimed to technically validate a new tropicalized hop cultivar, “Carnaval,” developed over six years of breeding and genetic improvement to adapt the crop to Brazilian conditions. A comparative analysis was conducted between Carnaval and two commercial varieties, Comet and Cascade, evaluating morphological traits, production potential, phytochemical composition via GC-MS, and genotyping using 10,541 SNP markers obtained through dd-GBS on the Illumina® NextSeq™ 1000/2000 platform. Carnaval exhibited clear genotypic and phenotypic distinctiveness, with notable differences in cone, leaf, and branch morphology, as well as in alpha (14.0%) and beta (6.9%) acid content and essential oil concentration (0.61%), comprising over 35 components—mainly myrcene, α-humulene, selinene, and caryophyllene. Productivity reached 478 kg/ha, comparable to commercial cultivars. Genetic analyses revealed high diversity (HS = 0.35), and population structure showed Carnaval grouping with its female parent Zeus but with distinct genomic characteristics. These findings confirm Carnaval’s unique identity and its adaptation to tropical conditions, making it a promising alternative for domestic hop production in Brazil.

## 1. Introduction

Keep the introduction short, but include (i) a brief statement of the problem that justifies doing the work, or the hypothesis on which it is based; (ii) the findings of others that will be further developed or challenged; and (iii) an explanation of the general approach and objectives. This last part may indicate the means by which the question was examined, especially if the methods are new.

Hops (*Humulus lupulus* L.) belong to the *Cannabaeceae* family and have a composition rich in bioactive compounds which are responsible for giving beer its characteristic flavor and aroma. These compounds can also be used for various other purposes such as in the formulation of drugs, supplements and cosmetics (Rutnik, et al., 2022).

The global hops market generates almost US$ 8 billion annually, with this number projected to almost double in the next ten years. Beer is the most consumed alcoholic beverage in the world and the third overall, behind only water and tea (Mordor Intelligence, 2024).

Hop production has been undergoing changes driven by the growth in the consumption of local and ‘craft’ beers, which has been increasing the demand for greater aromatic and flavor diversity and bitterness. When compared to traditional recipes, artisanal recipes use a greater concentration and variety of hops (González-Salitre et al., 2023). Thus, in addition to increasing yield, the hop production chain has sought to increase the diversity of materials offered.

The chemical composition of the crop is influenced by genetic factors and soil and climate conditions, thus, the concept of terroir has been applied to the hops used for beer. A study by Van Holle et al. (2021), demonstrated that the commercial cultivar Amarillo cultivated in Germany, the USA, and Belgium present distinct sensory attributes, despite the samples evaluated being in different temperate climate regions and having used the same genotype.

In this sense, the expansion of cultivation in different climatic regions can contribute to increasing the production and aroma and flavor diversity of the crop. Tropical regions were historically considered unsuitable for hop-cultivation. In 2019, there was no relevant hop production outside the temperate climate (Kubeš, 2022). However, recent studies have shown that crop productivity is more correlated with the long photoperiod then the actual temperature and climate where it is grown. (Bauerle, 2019). Thus, the adaptation of the crop to a tropical climate can occur by supplementing light hours or by developing cultivars adapted to tropical conditions (MAPA, 2020).

In this context, the present study presents the new cultivar “Carnaval” developed in Brazil from over 10 thousand crosses between commercial and experimental cultivars, which demonstrated adaptation to tropical conditions and may contribute to the expansion of the crop.

## 2. Materials and Methods

### 2.1 Field experiment

The new hybrid “Carnaval” developed by Hops Brasil Cultivo e Comércio de Lúpulo Ltda. was evaluated. It was previously selected in preliminary studies from over 10 thousand crosses and preliminary field tests with crosses that were selected for advanced testing due to their presenting better adaptation to water stress, good vigor, productivity, disease and pest resistance and desirable chemical characteristics in the preliminary tests. Twenty one hybrids were selected for advanced field testing.

The test was conducted in a commercial cultivation area and had total twenty three seperate hops cultivars, which included twenty one pre-selected hybrids and the two commercial cultivars.

A randomized block design with four replications was adopted, with the experimental plot represented by 4 groups each of 5 plants with spacing of 1.00 m between plants and 3.5 m between rows, corresponding to 20 plants distributed in 70 m2 per cultivar, totaling 460 plants and 1470 m2 of experimental area.

### 2.2 The experiment included one agricultural cycle of the crop

The phenological scale proposed by BBCH (Rossbauer, 1995) was used to monitor the evolution of the plants. When they reached stage 7 of cone maturation, an evaluation was performed on 10 plants of each cultivar by phenotypical analysis of the following items: stem pigmentation and diameter, plant height, shape of the productive part of the plant, counting the number of cones per branch, height of the first lateral branch containing cone production; leaf count per branch; number of cones per plant, fresh cone production, cone mass per plant, cone length and width, size and shape of the cone rachis and petals.

Each parameter was analyzed separately in analysis of variance (ANOVA), and data with normal distribution and were submitted to the Tukey test (p ≤ 0.05). Multivariate analysis was also performed between the DHE parameters and chemical analyses of the new hybrids and commercial materials. For this purpose, the Euclidean distance was calculated and a graph of dissimilarity between the genetic materials was plotted. The Mini Tab Statistical Software was used.

### 2.3 Chemical characterization

The 21 pre-selected hybrids and the Cascade and Comet cultivars grown in Brazil and the USA were evaluated for their chemical composition in duplicate. Each sample consisted of 150 g of fresh cones. To prepare each sample, 3 plants were randomly selected; approximately 50 g of flowers from female plants (cones) with golden-colored resins and an intense lupulin aroma were collected from each plant. The samples were vacuum-packed and kept frozen until the time of analysis.

The cones were analyzed for total alpha acid content using the spectrophotometric method described by the American Society of Brewing Chemists, method ASBC 6A, at the Hops Brasil laboratory.

The essential oil was extracted from the cones of each cultivar by hydrodistillation in a Clevenger apparatus at the Hops Brasil laboratory. The essential oil samples were sent for aromatic profile analysis by gas chromatography coupled to mass spectrometry (GC-MS) at the Agronomic Institute (IAC), Campinas, SP. The analyses were performed on a Thermo Scientific gas chromatograph (model TRACE 1300 Series GC) coupled to a mass spectrometer (model ISQ 7000) and a Triplus RSH automatic injector. The injector was maintained at 220 °C with a carrier gas flow rate (helium, 99.9999% purity) divided in a ratio of 1:20. The mass spectrometer (MS) operated in full scan mode, by electron impact (70 eV) and acquisition range of 40 to 450 m/z. The interface temperature was maintained at 220°C and the MS transfer line at 250°C (Adams, 2017).

The separation of the substances was performed in an Rtx-5 MS capillary column (30 m x 0.25 mm x 0.25 μm), with a carrier gas flow rate of 1.0 mL.min-1 in the following temperature program: 60°C - 240°C, 3°C.min-1. The Chromeleon® software (Thermo Scientific-Waltham, MA, USA) was used for data acquisition and processing.

### 2.4 Genetic sequencing

Cultivars were identified by comparative analysis of mass spectra with the National Institute of Standards and Technology (NIST 14); Flavour & Fragrance Natural & Synthetic Compounds (FFNSC3) libraries and linear retention indices of cultivars (IRL) with the literature (Adams, 2017). Linear retention indices were obtained by injecting a series of n-alkanes (C9-C24, Sigma-Aldrich, 99%) under the same chromatographic conditions as the samples, applying the Van den Dool and Kratz equation (1963).

DNA was extracted from young leaves using the protocol based on the CTAB 3% reagent (cetyltrimethylammonium bromide) (Doyle and Doyle 1990). The samples were quantified on a 1% (w/v) agarose gel and standardized to a concentration of 30 ng/µl for the preparation of the GBS-based genomic library (Genotyping-by-Sequencing) following the double digestion protocol described by Poland et al. (2012). First, the genomic DNA of each individual was digested by two restriction enzymes (Pst I and Mse I). These were chosen after testing considering the genome size and the amount of repetitive sequences. The digested fragments of each sample were ligated to individual adapters (barcodes) (Sigma-Aldrich, St. Louis, MO, USA) and to specific adapters for Illumina technology (Illumina, CS, Pro). The restriction-ligation products were then pooled into a single pool (multiplex), which was purified using the QIAquick PCR Purification Kit (QIAGEN®), and amplified by PCR to enrich the fragments with both restriction sites. After this step, the samples were again purified, validated and quantified on a Qubit V3.0 Fluoremeter device (Thermo Fisher Scientific, Carlsbad, CA, USA), to be subjected to sequencing on the Illumina platform (Illumina, Inc.).

## 3. Results and Discussion

The evaluation of development, production parameters, chemical and genetic analyzes made it possible to distinguish six new advanced hybrids in relation to commercial cultivars.

The morphological characteristics of the two commercial materials (Cascade and Comet) and of five new hybrids: Carnaval, 2199/201, 2123/241, 2119/211, 2002/125 and 2001/103, allowed us to assess that the new hybrids differ from those commercially available. It is noted that most of the parameters presented differences between the cultivars, indicating homogeneity between plants of the same genetic material and difference between the hybrids.

The morphological analysis of the new cultivar Carnaval is illustrated in Figure 1. The following parameters present significant differences in relation to at least one of the commercial cultivars, reddish pigmentation of the stem, medium leaf size, medium intensity green leaf coloration, long lateral shoot length with dense foliage.

**Figure 1.**
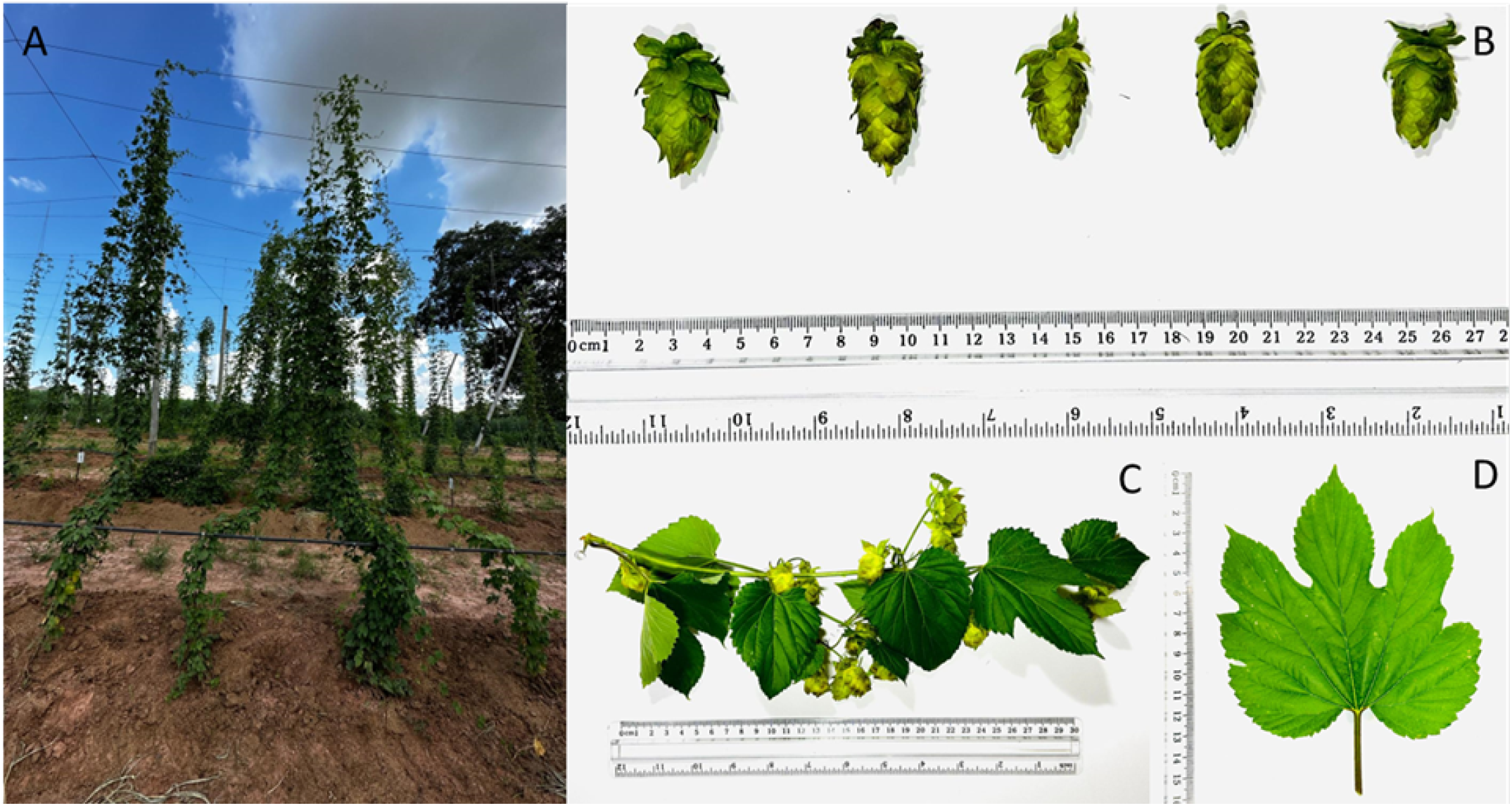
New hop cultivar “CARNAVAL” HB2001/114 in field test (A – Plant at stage 7, BBCH scale; B – Cones and bracts; C – Lateral branch; D – Leaf).

The plant size of all the materials evaluated was considered normal and with a height of 6 m, the same for all and equal to the maximum height of the support structure.

The new hybrids presented a later grow cycle in relation to the commercial cultivars, that is, the number of days between planting and harvesting was greater for most hybrids in comparison with Comet and Cascade, both early with a cycle of 119 days, with Carnaval presenting 125 days, reaching 148 days for 2002/125 and with the exception of 2119/211 which presented the same cycle duration of 119 days as the commercial ones.

The materials 2123/241, 2119/211 and 2001/103 were considered similar to each other in relation to the others regarding the mass of fresh cones per plant and presented the highest results for productivity estimation, with 786.3 kg.ha-1, 1057.3 kg.ha-1 and 862.8 kg.ha-1, respectively. The three hybrids also stood out in relation to the number of cones per branch. The commercial materials (Comet and Cascade) and the hybrids 2002/125 and Carnaval presented similar results to each other and lower than the others for productivity, reaching 431.4 kg.ha-1 for Cascade. Regarding the quality of the cones produced, the hybrids 2123/241, 2119/211, 2001/103 and Carnaval presented similar results for alpha and beta acids and higher than those presented by Comet and 2002/125, reaching 14% alpha acid and 6.8% beta acid for Carnaval.

The new cultivar also stood out for its essential oil content, of approximately 0.61%, considered similar to the commercial cultivar Cascade. The hybrids 2119/211 and 2001/103 presented intermediate values that were considered similar to the commercial cultivar Cascade, reaching 0.62% for 2119/211. And the hybrid 2123/241 presented the lowest content, of 0.42%. The high number of parameters analyzed made it difficult to perform a global analysis of the new hybrids. Therefore, we opted for multivariate dissimilarity analysis to measure the distance between the different materials studied. Figure 2 shows the dendrogram of the Euclidean distances between the commercial cultivars and new hybrids in relation to the morphological criteria. In it, it can be noted that the commercial materials Cascade and Comet were considered more similar to each other in relation to the rest, with a distance of 3.48%. Carnaval, on the other hand, presented a greater distance in relation to the others, being closer to 2002/125, with a distance of 3.48% between them. The extracted essential oil samples were sent for analysis of the profile of aromatic compounds. In the Carnival samples, 35 substances were found in the samples, among which more than 80% of the chemical component proportional to the totality of each sample corresponded to five substances, namely, caryophyllene, myrcene, linanol, selinene and humulene. The high number of compounds found is similar to that found in the literature on the essential oil of the crop (Santagostini et al., 2020; Yan et al., 2019; Bocquet et al., 2018). However, the aromatic characteristics are related to the profile of components, including minor ones such as farnesene, geraniol, β-pinene. Thus, the comparison between the genetic materials was carried out in relation to the total components and their proportions, as illustrated in Figure 3. In which, it can be seen that the commercial cultivars that were grown in Brazil and the USA, despite being clones and, therefore, having the same genotype, were not classified as similar. Comet USA was considered the most distinct among all the essential oils evaluated, while Comet BR was considered most similar to 2123/241 and Cascade USA. This corroborates the initial hypothesis that soil and climate conditions influence the composition of the essential oil, reinforcing the concept of terroir similar to that used for grapes used for wine production. However, other variables were raised during the data analysis, such as whether there are significant differences in the composition of the same genotype cultivated in the same location in different harvests; whether the small scale of the experiment carried out in Brazil may have influenced its management and, consequently, the composition of the oils. Therefore, it is important to carry out new studies on the influence of environmental factors on the composition of the phytochemical profile of hops.

**Figure 2.**
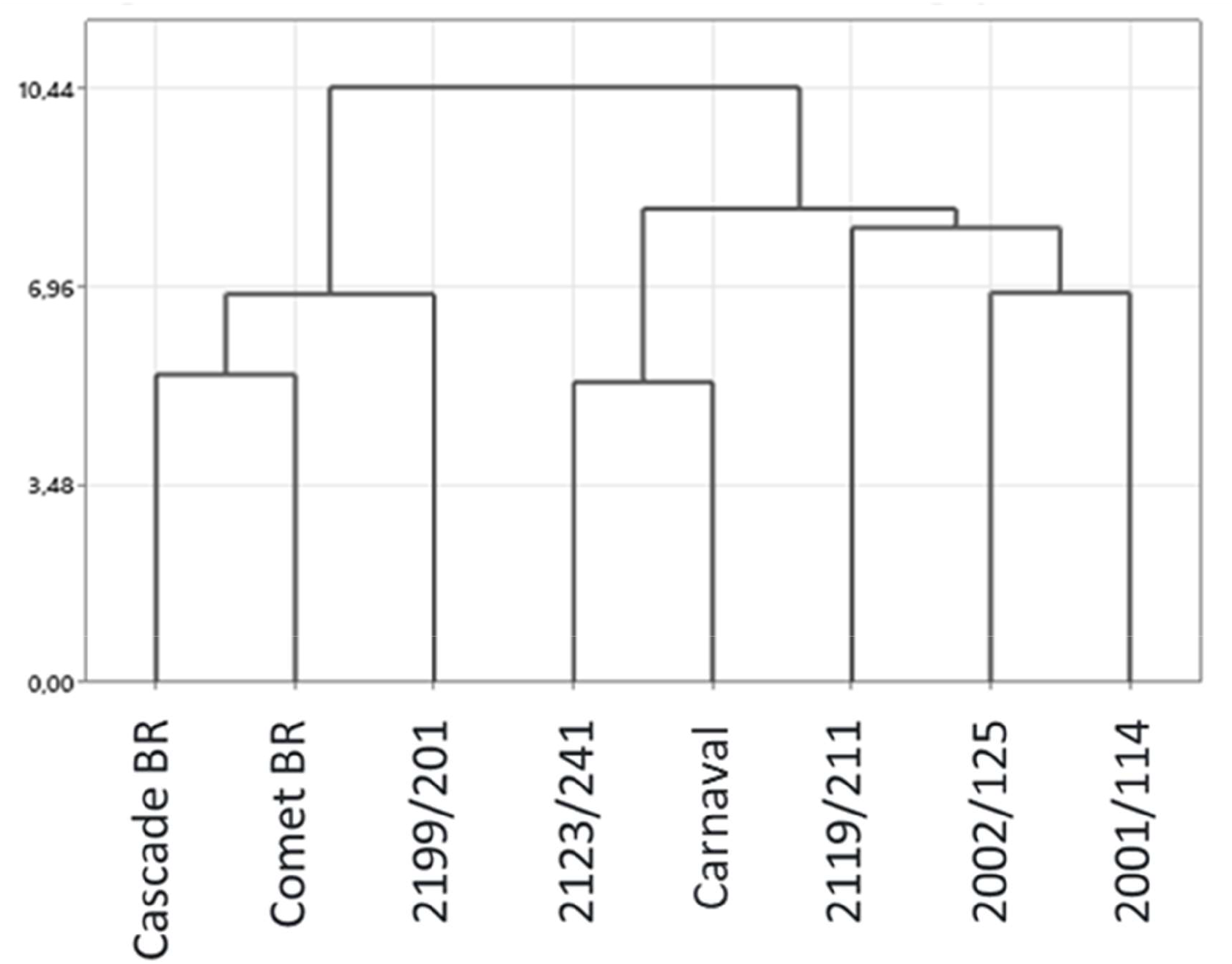
Study of dissimilarity of morphological criteria between commercial hop cultivars Cascade and Comet and new hybrids HB2001/114, HB2199/201, HB2123/241, HB2119/211, HB2002/125 e HB2001/114.

**Figure 3.**
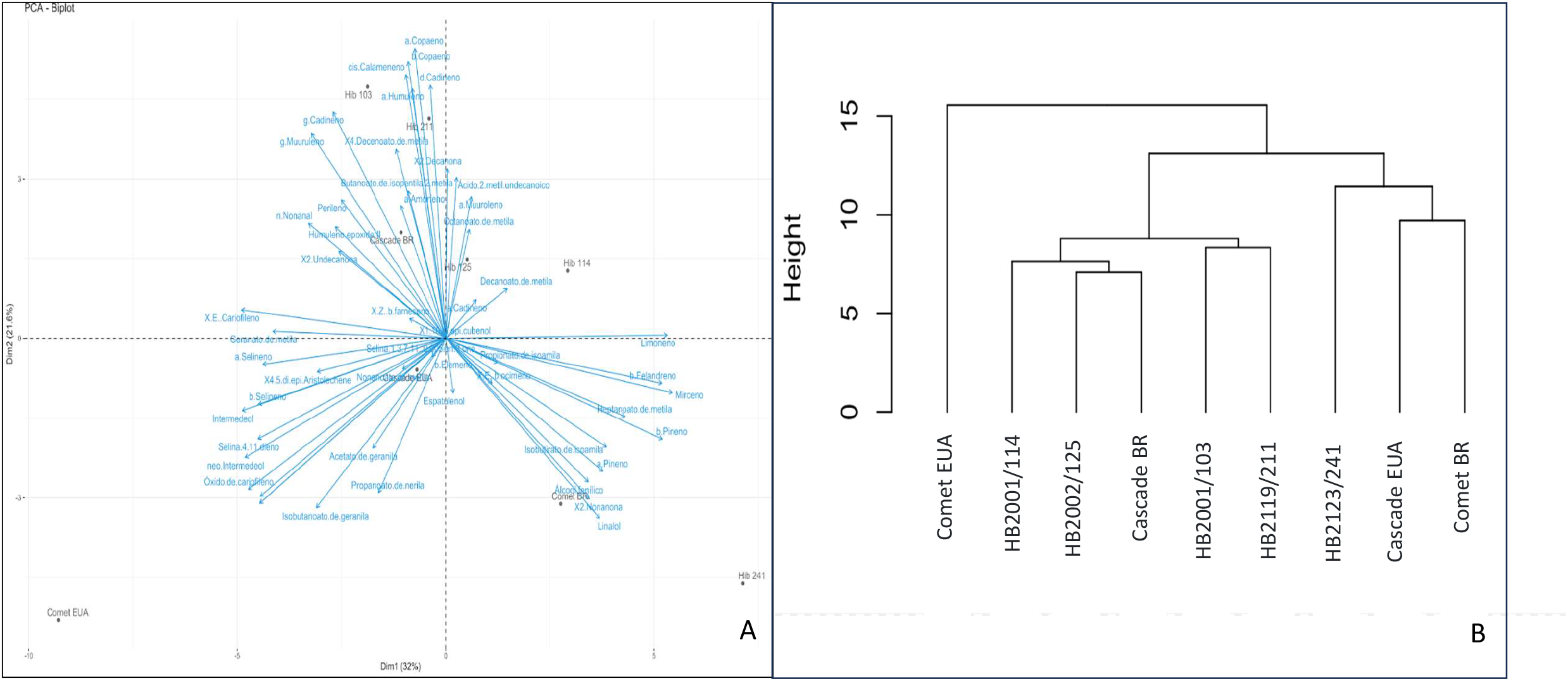
PCA analysis of chemical composition of hop essential oil (A - PCA per cultivar; B - Dendrogram per cultivar).

In addition, it was also found that the new hybrids produced essential oils with a distinct phytochemical profile from each other and in relation to commercial materials, also reinforcing the hypothesis that they are distinct from commercial cultivars and, therefore, are eligible for protection.

The hybrids and commercial cultivars were also subjected to genetic analysis using SNP (single nucleotide polymorphism) markers by the dd-GBS (double digest genotyping by sequencing) approach on the Illumina® NextSeq™ 1000/2000 next-generation sequencing (NGS) platform. A total of 166,388 loci were genotyped in 50 individuals from the Hops Brasil breeding program. The average coverage per sample was 36.33 × (SD = 20.7×, min = 11.8×, max = 72.7×). After filtering, the library resulted in 10,541 high-quality SNP markers. For the population analyses, 14 genotypes that are part of the commercial hybrids of interest were selected. The identification of a high number of SNP markers found in the samples is similar to the study conducted by Mafakhero (2020) on the assessment of genetic diversity of wild hops.

According to the 166,388 putatively neutral loci, the genotypes of Cascade x 101/19058 D, Comet, Hib 2002/125, Hib 2006-141 C, Hallertauer Magnum, presented the highest genomic diversity, estimated by the observed heterozygosity (Ho = 0. 37; 0.37; The HB2001/114 and Cascade genotypes also presented the highest gene diversity (HS = 0.35 and 0.306) and allelic richness proportion (AR). Regarding the number of alleles (A), the individuals with the highest values were Cascade x 101 19058 D, Hib HB2119/211 and Zeus (13,424, 13,259, 13,242 respectively). All of these genotypes are promising as future parents for the creation of tropicalized hop varieties. The hybrid HB2123/241 presented the lowest HO estimate (0.156). Regarding the proportion of AR, the lowest value was observed in male Rocky A (1.15).

Based on Rogers’ genetic distance, the Fst (fixation index) values between the genotypes defined two related groupings (Figure 4). The first grouping is made up of HB2002/125, Nugget, Hallertauer Magnum, Cascade, 19112 B, 2006-141 C, Comet and Hib 211. The second grouping is made up of Cascade x 101 19058 D, HB2123/241, HB2001/114, Zeus and Brewers Gold. These groups are differentiated from each other, as they occupy different quadrants within the DAPC (Figure 4.A) as in the dendrogram (Figure 4.B). The most distant varieties were Rocky A and HB 2119/211.

**Figure 4.**
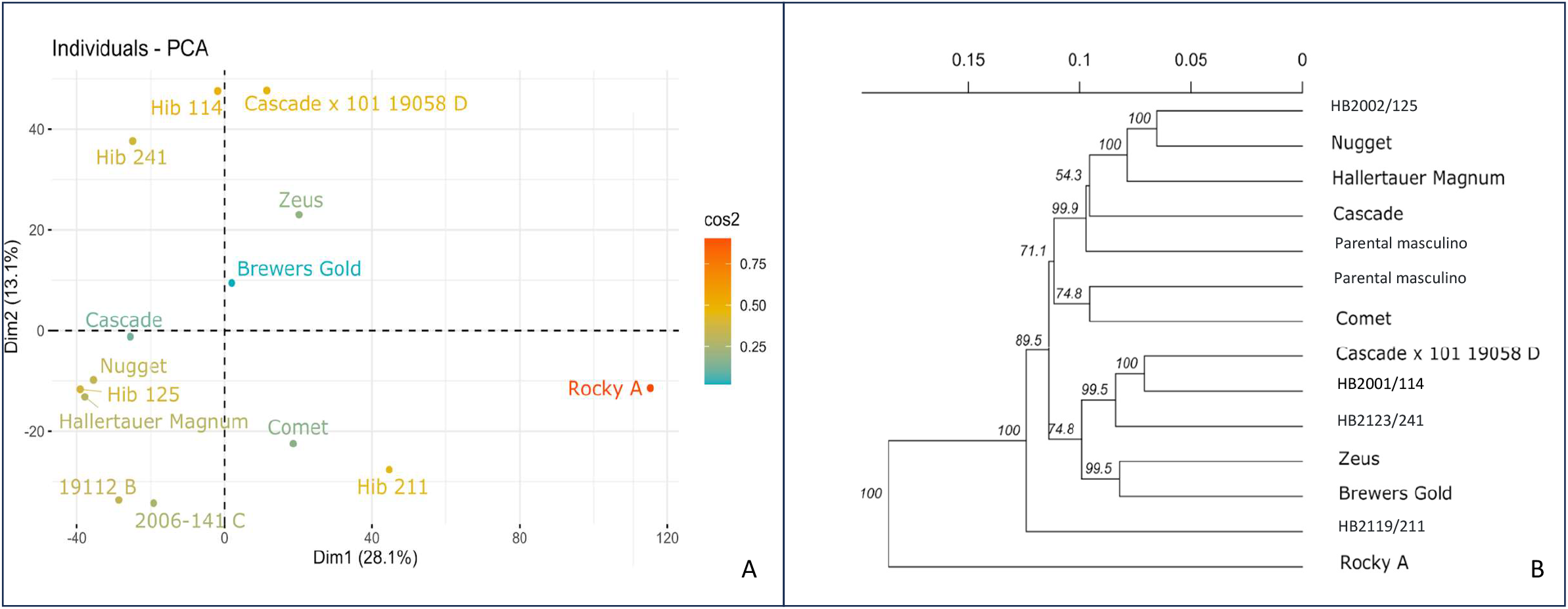
PCA analysis of hop molecular markers (A - PCA per cultivar; B - Dendrogram per cultivar).

To understand the genetic composition of the evaluated varieties, a structuring analysis was carried out. Like the Structure approach, the sMNF method assumes that population genomic data is the result of a mixture of K unknown parental lineages and estimates ancestry coefficients from multilocus genotypes. From K = 1 to 14 ancestral populations were tested, due to the 14 genotypes of interest, with 10,000 interactions, ten replications for each K value. To visualize the results of the simulations, the cross-entropy criterion was used. This analysis was carried out with the LEA package (Frichot & François, 2015) for the R 4.2.1 platform (R Development Core Team, 2022). Two ancestral parental lines were found in the 14 genotypes evaluated, thus indicating that the Hops Brasil germplasm bank has 2 different genetic clusters (Figure 5). This result demonstrates a close relationship between approximately 50% of the accessions, with the remainder discreetly moving away in a single direction, giving an idea of hop domestication. This analysis reinforces the difference between the genomic composition of the hybrids and the parents, showing that the varieties generated in the breeding program have their own brands that promote adaptation to Brazilian tropical conditions.

**Figure 5.**
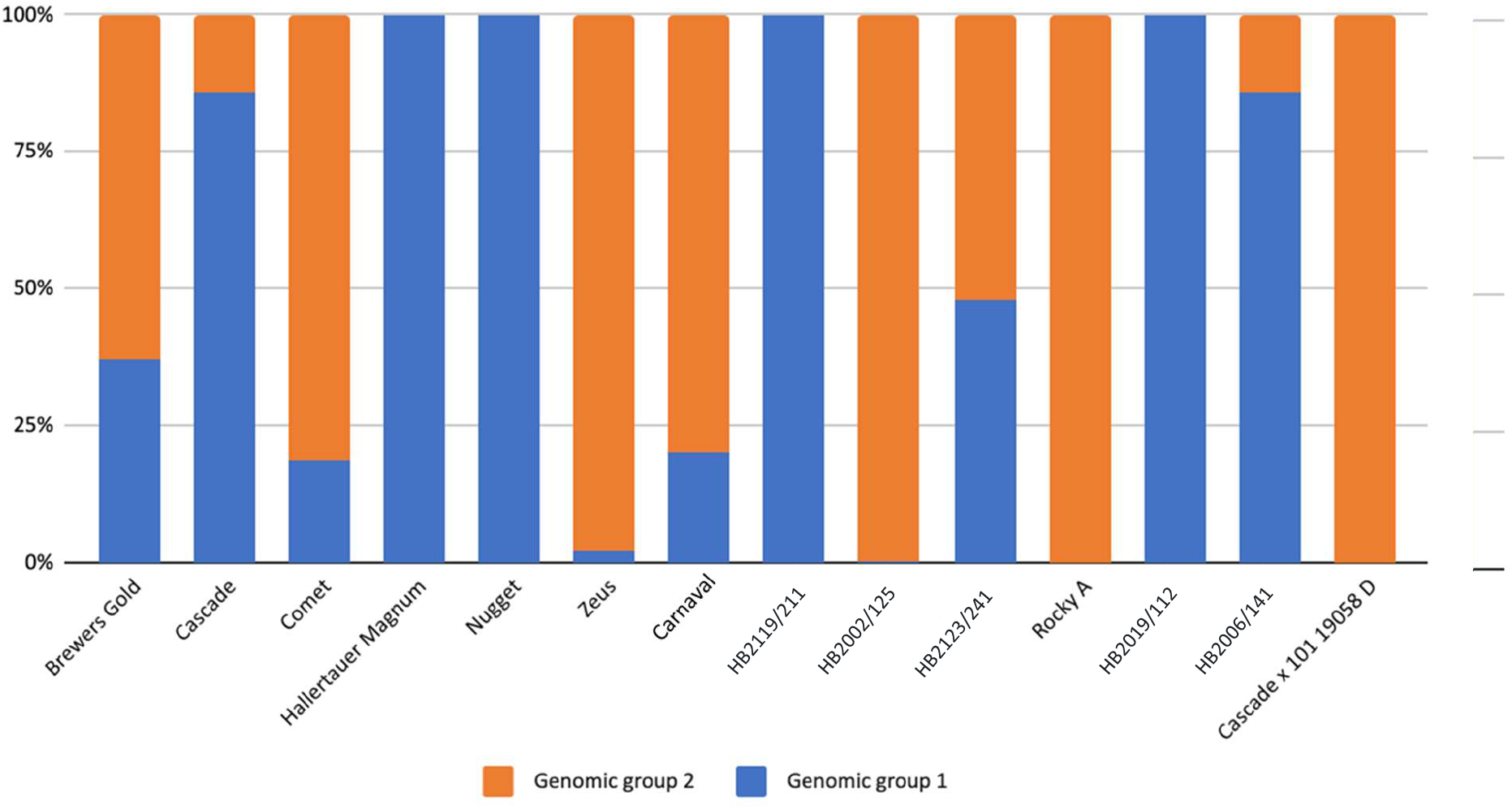
Genetic structuring of genotyped hop varieties.

The results strongly support that the efforts of the germplasm bank and hybrids generated in the Hops Brasil breeding program have brought together a diverse hop genetic base. The breeding program has highly heterozygous individuals generated from an extensive prospection of the hop gene pool. The hybrids evaluated showed a wide genetic differentiation within the collection, so that the geographic origin of the parents does not necessarily correspond to genetic kinship, thus showing the tropicalization of the culture in Brazil.

Thus, the data presented corroborate the initial hypothesis that the new hybrids differ from commercial cultivars, with emphasis on HB2001/114, HB2119/211 and HB2123/241, since they presented significant differences in most of the morphological criteria required for the application for cultivar protection; they demonstrated good cone productivity (female inflorescences), reaching 1057 kg.ha-1 for HB2119/211; the new materials presented values compatible with commercial hops in relation to the aromatic components alpha and beta acids, reaching 14% and 6.9% for α and β-acids respectively and 0.61% oil content for HB2001/114; thus, they demonstrated differentiated phytochemical profiles of essential oil; as well as distinguishing themselves from commercial and parental cultivars in relation to the genetic components evaluated. In this way, the study contributed to the advancement in the development of new hop cultivars, through a staggered and predictable production process of seedlings with genetic, phytosanitary and organoleptic guarantees, which will help in the popularization and dissemination of the crop in the country. With the potential to become a diversification option for small producers, since the crop has high added value, is little explored in the country and can be produced in small areas.

## 4. Acknowledgments

The authors thank FAPESP 2022/00356-9 for the financial support that made this study possible.

